# RAN translation of C9orf72-related dipeptide repeat proteins in zebrafish recapitulates hallmarks of amyotrophic lateral sclerosis and identifies hypothermia as a therapeutic strategy

**DOI:** 10.1101/2024.01.17.576077

**Authors:** David J. Burrows, Alexander McGown, Olfat Abduljabbar, Lydia M. Castelli, Pamela J. Shaw, Guillaume M. Hautbergue, Tennore M. Ramesh

## Abstract

**Objective:** Hexanucleotide repeat expansions in the *C9orf72* gene are the most common genetic cause of amyotrophic lateral sclerosis (ALS) and frontotemporal dementia (FTD). A large body of evidence implicates DPRs as one of the main drivers of neuronal injury in cell and animal models.

**Methods:** A pure repeat-associated non-AUG (RAN) translation zebrafish model of C9orf72-ALS/FTD was generated. Embryonic and adult transgenic zebrafish lysates were investigated for the presence of RAN-translated DPR species and adult-onset motor deficits. Using C9orf72 cell models as well as embryonic C9orf72-ALS/FTD zebrafish, hypothermic-therapeutic temperature management (TTM) was explored as a potential therapeutic option for C9orf72-ALS/FTD.

**Results:** Here we describe a pure RAN translation zebrafish model of C9orf72-ALS/FTD that exhibits significant RAN-translated DPR pathology and progressive motor decline. We further demonstrate that hypothermic-TTM results in a profound reduction in DPR species in C9orf72-ALS/FTD cell models as well as embryonic C9orf72-ALS/FTD zebrafish.

**Interpretation:** The transgenic model detailed in this paper provides a medium throughput *in vivo* research tool to further investigate the role of RAN-translation in C9orf72-ALS/FTD and further understand the mechanisms that underpin neuroprotective strategies. Hypothermic-TTM presents a viable therapeutic avenue to explore in the context of C9orf72-ALS/FTD.

## INTRODUCTION

Amyotrophic lateral sclerosis (ALS) is a fatal neurodegenerative disease that is characterized by the progressive degeneration of both upper motor neurons in the motor cortex and lower motor neurons in the spinal cord. Disease onset begins with skeletal muscle degeneration and progresses to eventual respiratory failure and death within 2-4 years of symptom onset^1^. The majority of ALS cases are sporadic (sALS), which account for approximately 90-95% of cases. The remaining 5-10% of cases have familial history (fALS) and a strong inherited link^2^. The *C9orf72* gene is the most common identified gene associated with ALS pathogenesis. Following a landmark paper published in 2011, it was identified that polymorphic hexanucleotide (GGGGCC)n repeat expansions (HREs), located in the first intron, between exons 1a and 1b of the *C9orf72* gene, are the most common genetic cause of ALS accounting for approximately 40% of fALS or 10% of ALS cases overall^3^. HREs in the *C9orf72* gene are also the most frequent cause of familial frontotemporal dementia (FTD)^4^. FTD encompasses a spectrum of neurocognitive disorders which present clinically with cognitive and/or language impairment, as a result of degeneration in the frontal and/or temporal cortical lobes. Because FTD and ALS share genetic, clinical and pathogenic mechanisms that lead to disease initiation and progression, C9orf72-ALS/FTD is used as a collective term for C9orf72-associated diseases where patients present with ALS, FTD or both symptoms^5^.

Although neurologically healthy people have ≤ 30 hexanucleotide repeats in the *C9orf72* gene, larger expansions of hundreds to thousands are observed in C9orf72-ALS/FTD patients^6^. The pathological pathways associated with C9orf72-ALS/FTD involve both loss-of-function and gain-of-function mechanisms. These have been broadly categorized into three potentially overlapping pathophysiological mechanisms and include: (1) loss of function of the C9orf72 protein due to decreased transcription of the *C9orf72* gene; (2) toxic gain of function from sense and antisense repeat RNA sequestering or altering RNA-binding protein function and; (3) the production of toxic dipeptide repeat (DPR) proteins via repeat associated non-AUG (RAN) translation (for review see^7^). The mechanisms of RAN translation remain poorly characterized, however it involves highly structured RNA and occurs in all three reading frames in absence of the canonical AUG initiation codon via the near cognate initiation codons CUG or GUG (for review see^8^).

The development of novel animal models of C9orf72-ALS/FTD has given pre-clinical researchers the capacity to interrogate and gain further insights into the complex pathological mechanisms that contribute towards ALS pathogenesis and progression. In recent years, zebrafish (*Danio rerio*) have emerged as a useful pre-clinical tool to study gene function in the context of human disease, including C9orf72-ALS/FTD^9, 10^. Zebrafish models offer a low-cost and higher throughput alternative when compared to traditional rodent models. Their rapid external development, translucent embryos and amenability to genetic manipulation facilitate the opportunity for high-to medium-throughput therapeutic screening. The zebrafish genome shares 70% homology with human genes and over 80% of human disease-associated genes have a zebrafish gene counterpart^11^, highlighting their utility as a pre-clinical model system for further understanding vertebrate gene function in health and disease.

Hypothermic-therapeutic temperature management (TTM), or therapeutic hypothermia, has been identified as an alternative, non-invasive therapeutic avenue for neurodegenerative disease. A protective role of hypothermic-TTM has been reported in a number of acute neurological and cardiovascular diseases, including stroke, traumatic brain injury (TBI), hypoxic-ischemic encephalopathy and cardiac arrest^12^. Through these studies, clinical treatment protocols have been refined and the mechanisms that underpin this neuroprotection have been explored. Although these pathways have not yet been fully elucidated, profound effects on cellular metabolism, production of cold-shock proteins/hormones^13^, free radical generation^14^, synaptic plasticity^15^ and BBB disruption^16^ have all been identified as possible contributory mechanisms. The potential therapeutic effects of hypothermic-TTM has not yet been explored in the context of C9orf72-ALS/FTD.

This study aims to build on this work by exploring hypothermic-TTM in the context of C9orf72-ALS/FTD by utilising cellular models of C9orf72-ALS/FTD and a zebrafish model of C9orf72-ALS/FTD. We first describe a novel pure RAN translation zebrafish model of C9orf72-ALS/FTD that exhibits significant RAN-translated DPR pathology and progressive motor decline, recapitulating clinical tissue pathology and clinical phenotypes. Further, we build on this previous work by demonstrating a profound reduction in DPR species following the implementation of hypothermic-TTM in C9orf72 cellular models as well as embryonic C9orf72-ALS/FTD zebrafish.

## MATERIALS AND METHODS

### Transgene constructs

RAN translation plasmids expressing 45 repeats of the *C9orf72* sense hexanucleotide GGGGCC (G4C2) and 43 of the antisense hexanucleotide CCCCGG (C4G2) with V5 tags in all three open reading frames^17^ were used to subclone the repeat expression cassettes into zebrafish plasmids under a ubiquitin (*Ubi*) promoter. Sanger sequencing showed that 45 sense and 39 antisense repeats were subcloned into the zebrafish plasmids co-expressing a red fluorescent protein (DsRed) under the heat shock protein *Hsp70* stress response promoter. Embryos were injected with these constructs or a plasmid containing two G4C2 repeats (2mer control) to generate transgenic zebrafish lines as previously described^10^. Potential GUG start codons are located 12 nucleotides and 17 nucleotides in the 0 and +1 reading frames upstream of the 45 sense repeats. Potential GUG start codons are present 10 nucleotides and 5 nucleotides in the 0 and −1 reading frames upstream of the 39 antisense repeats. A diagrammatic representation of the constructs is outlined in Figure 1. Zebrafish lines were maintained using established practices^18^.

**Figure 1.**
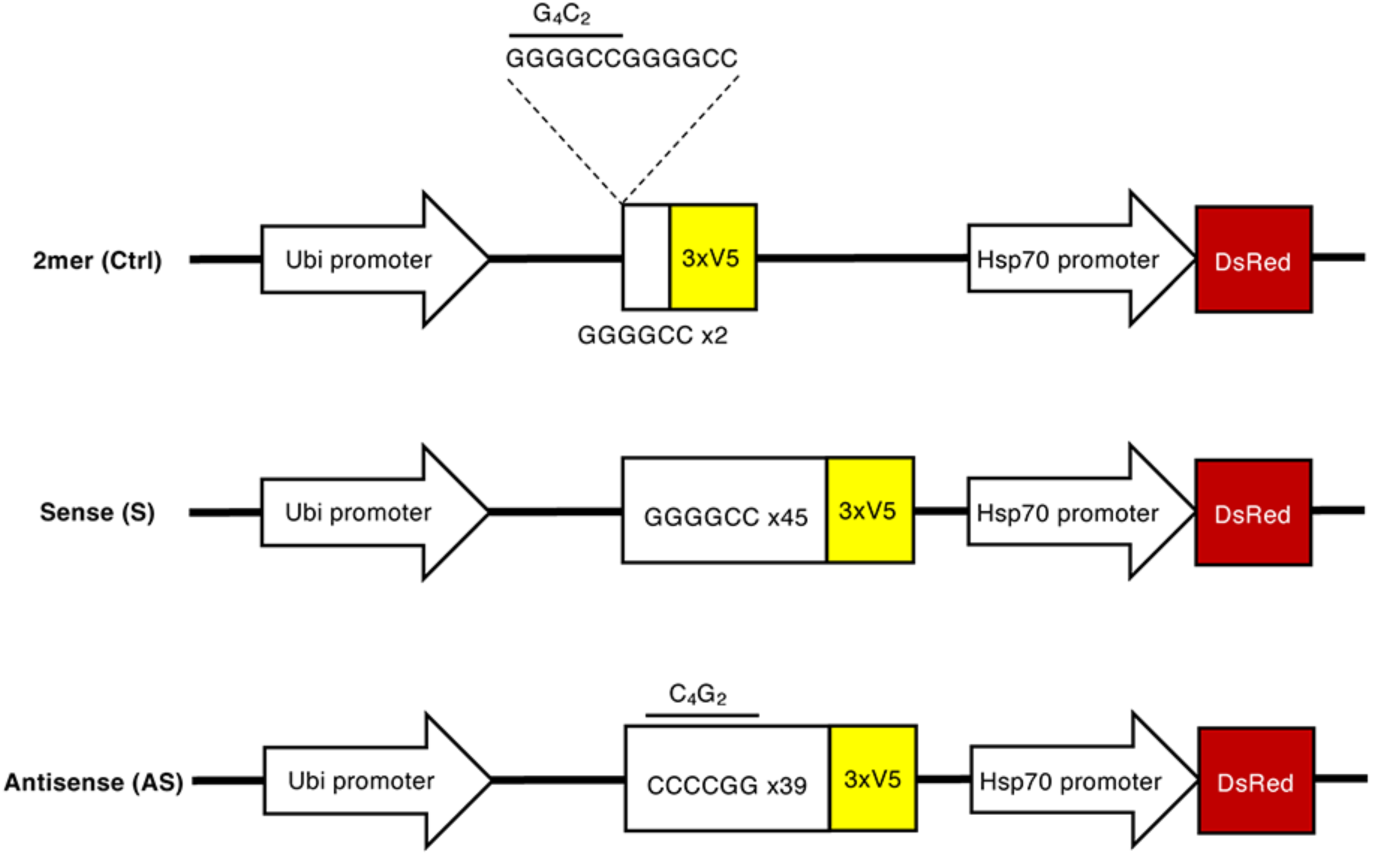
Schematic of transgene used to create pure repeat RAN transgenic C9orf72-ALS/FTD zebrafish lines. The 2mer control line contains 2 repeats of GGGGCC (G4C2), the sense (S) line contains 45 repeats of G4C2 and the antisense (AS) line contains 39 repeats of CCCGG (C4G2). A V5 epitope tag is in all three frames following the repeats (3xV5) and transgenes also carry the hsp70-DsRed transgene. No ATG codon exists that could allow the translation of the repeats in all three frames.

### Western blotting

Whole zebrafish embryos were processed for western blotting following terminal anesthesia using MS-222 (Sigma). Laemmli buffer was added to whole anaesthetized zebrafish embryos in the ratio of 8 μl per embryo, homogenized using the Precellys 24 (Bertin Instruments) tissue homogenizer (2500 rpm, 2×20 secs) and then centrifuged at 13,000 g for 10 mins. Supernatant was collected and boiled at 95°C for 10 min prior to SDS-PAGE and immunoblotting Commercial mouse primary antibodies used were α-V5 (Sigma, R960), α-tubulin (Abcam, ab6046), as well as the rabbit primary antibodies α-DsRed (Clontech, 632-496), α-CIRBP (Proteintech, 10209-2-AP) and α-RBM3 (Proteintech, 14363-1-AP). All other DPR-specific rabbit antibodies targeting poly(GP), poly(GR) and poly(PR) were custom-generated (Eurogentec) and purified in-house using affinity membrane chromatography. Species-specific HRP-conjugated secondary antibodies were used and immunoblots imaged by chemiluminescence using LiCor Odyssey® Fc imaging system.

### Swim tunnel analysis

Zebrafish swimming ability was tested using a swim tunnel with an initial flow-rate of 6.6 cm/sec, increasing in 6.6 cm/sec increments every 5 min until the maximum flow rate of 39.5 cm/sec was achieved. Data were analysed as previously described^10^. C9orf72-ALS/FTD transgenic zebrafish may show delayed recovery, which manifests as a reduction in swimming activity, immediately following the swim tunnel assessment.

### Cell culture and transfection

Cells were maintained in a 37°C incubator with 5% CO_2_. HEK293T and HeLa cells were cultured in Dulbecco’s Modified Eagle Medium (Sigma), supplemented with 10% foetal bovine serum (FBS) (Gibco) and 5 U ml^−1^ Penicillin-streptomycin (Lonza). Neuro-2a (N2A) (ATCC) cells were cultured in Dulbecco’s Modified Eagle Medium (Sigma), supplemented with 10% FBS (Gibco), 5 U ml^−1^ Penicillin-streptomycin (Lonza) and 5 mM sodium pyruvate. HEK293T, HeLa and N2A cells were transfected with 700 ng of plasmid using 3.5 μg PEI/ml media and one tenth media volume of OptiMEM in a 24 well format. Approximately, 50,000 HEK293T cells, 50,000 HeLa cells and 75,000 N2A cells were seeded per well of the 24 well plate.

### Hypothermic treatment

#### In vitro

24 h post-transfection, cells were washed and complete media replaced. Plates were incubated for 48 h at 37°C (normothermia) or 32°C (hypothermia) before proteins were extracted 48 h post-treatment. Cells were washed in ice cold phosphate buffered saline (PBS) and subsequently lysed in ice cold lysis buffer (50 mM HEPES pH 7.5, 150 mM NaCl, 10% glycerol, 0.5% Triton X-100, 1 mM EDTA, protease inhibitor cocktail (Sigma), 1 mM DTT) for 10 min on ice. Extracts were then centrifuged at 17,000 g for 5 min at 4°C. Extracts were quantified using Bradford Reagent (BioRAD), resolved by SDS-PAGE, electroblotted onto nitrocellulose membrane and probed with the relevant primary antibodies.

### Zebrafish

Embryos were manually dechorionated and transgenic positive fish identified by DsRed expression at 2 days postfertilization (dpf). Only normally developed and healthy appearing embryos, with expected fluorescence patterns, were selected for screening. Embryos were incubated at 28°C (normothermia) or 22°C (hypothermia) for 2-5 dpf. The zebrafish larvae were then terminally anaesthetized using MS-222 (Sigma) and homogenized in the Precellys 24 Tissue Homogeniser in 1X RIPA solution (Merck-Millipore) before being centrifuged at 17 000 g for 15 min. Supernatant was collected for downstream analysis.

### MSD poly(GP) immunoassay

A poly(GP) Meso Scale Discovery (MSD^®^) enzyme-linked immunosorbent assay (ELISA) was established using a custom made rabbit α-GP antibody (Eurogentec) and based on previously described methods^19^. Briefly, 30 μl of poly(GP) antibody (4 μg ml^-1^) diluted in 1X tris-buffer saline (TBS) was added per well of a 96-well SECTOR plate (MSD) and incubated overnight at 4°C. Plates were washed three times in 1X TBS-Tween^®^-20 (0.1%) and blocked for 1 h shaking in 150 μl 3% milk-TBS-T solution (RT, 700 rpm). Plates were washed again in TBS-T and 50 μl GP_(7)_ standard (0-400 ng / mL) or zebrafish lysate was added per well and incubated for 2 h shaking (RT, 700 rpm). Plates were washed again three times in TBS-T and 50 μl MSD^®^ SULFO-TAG labelled streptavidin (1 μg ml^-1^) and biotinylated poly(GP) antibody (2 μg ml^-1^, Eurogentec) diluted in blocking solution was added per well and incubated for 2 h shaking (RT, 700 rpm). Wells were washed again, 150 μl 2X MSD^®^ read buffer-T added and plates imaged using the MESO SECTOR 2400 imager. A synthetic poly(GP) peptide containing seven glycine-proline repeats (GP7) was used as a standard curve to interpolate assay response values prior to normalization.

### Quantification and statistical analysis

Data were analysed by one-way ANOVA with Tukey’s post hoc test or two-way ANOVA with Sidak’s post hoc test for multiple comparisons, t-test or Kaplan Meier analysis as indicated in the appropriate figure legend. Significance is denoted as * P < 0.05, ** P < 0.01, *** P < 0.001 and **** P < 0.0001.

## RESULTS

### Embryonic and adult C9orf72-ALS/FTD zebrafish express multiple RAN-translated DPR species

Five DPR species are generated in C9orf72-mediated disease and have been shown to be toxic in a number of CNS *in vitro* and *in vivo* models^20^. Therefore, our first aim was to characterize the DPR species produced in each of the generated transgenic zebrafish lines. Western blot analysis of 5 dpf zebrafish embryo lysates was performed to determine the presence of RAN translated V5-tagged DPR species in three 2mer (Ctrl), five sense (S) and four antisense (AS) founder transgenic zebrafish lines. Western blot analysis showed varying levels of RAN-mediated V5 expression between the generated lines (Figure 2). These ranged from high, moderate to low expression levels. No V5 expression was detected in any of the three transgenic founder 2mer (Ctrl) lines generated. In comparison with the other transgenic founder lines generated, sense line S5 and antisense line AS1 exhibited the highest expression of V5-tagged DPRs that correlated with the DsRed stress response (Figure 2). Therefore, sense line S5 and antisense line AS1 were chosen for further characterisation of DPR species in embryonic and adult tissue lysates.

**Figure 2.**
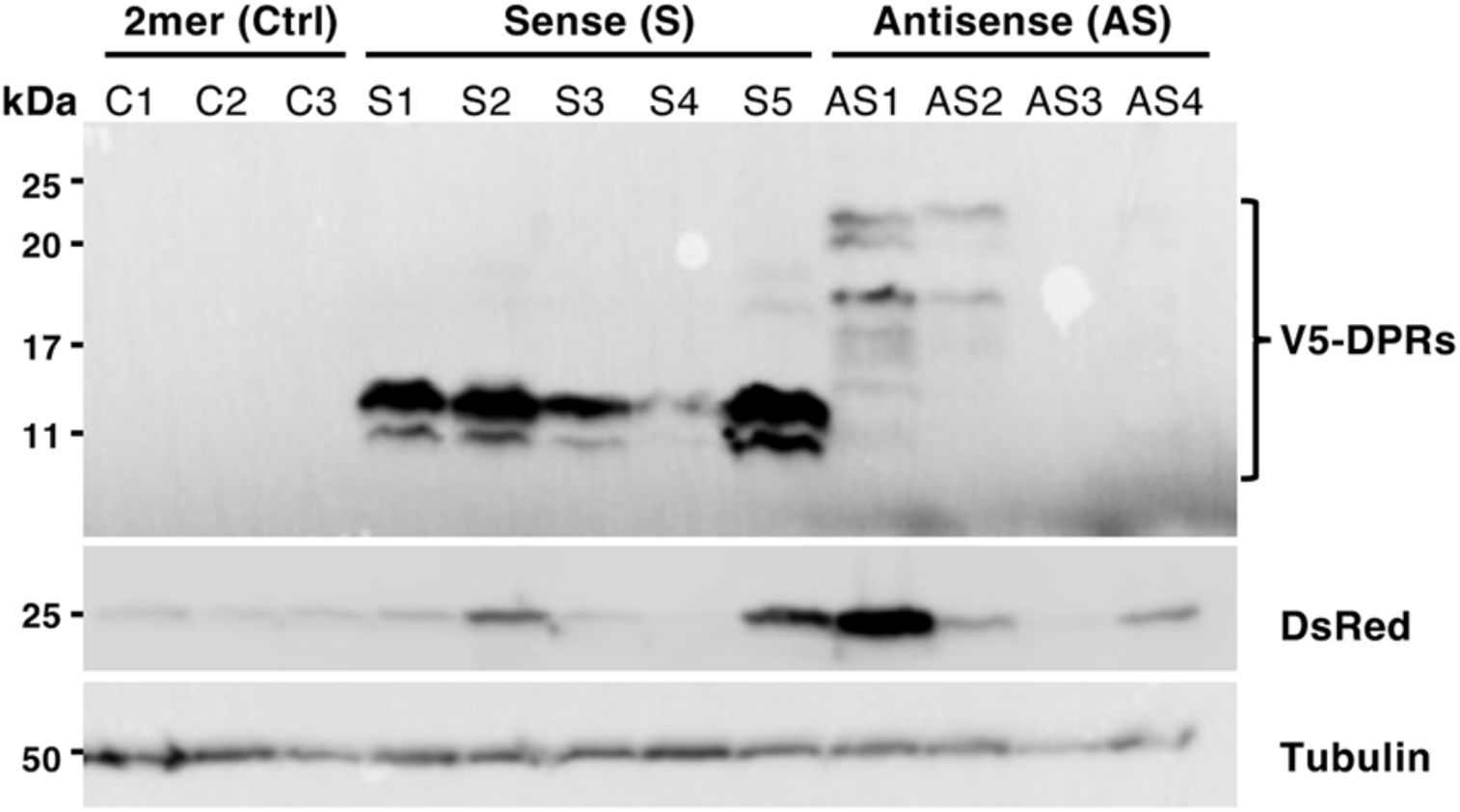
Western blot of RAN-translated products in transgenic 2mer (Ctrl), sense (S) and antisense (AS) 5 dpf embryonic zebrafish. Three 2mer (Ctrl), five sense (S) and four antisense (AS) generated transgenic lines were probed with anti-V5, anti-DsRed and anti-tubulin (loading control) antibodies.

V5 immunostaining demonstrated the production of multiple V5-tagged DPR species at different molecular weights in both sense line S5 and antisense line AS1 (Figures 3A-B). The sense line S5 expresses poly(GP) DPRs as well as poly(GR) DPRs, albeit to a lesser extent (Figure 3A). Compared to the sense line, the antisense AS1 line exhibits high expression of both poly(GP) and poly(PR) DPR species (Figure 3B).

**Figure 3.**
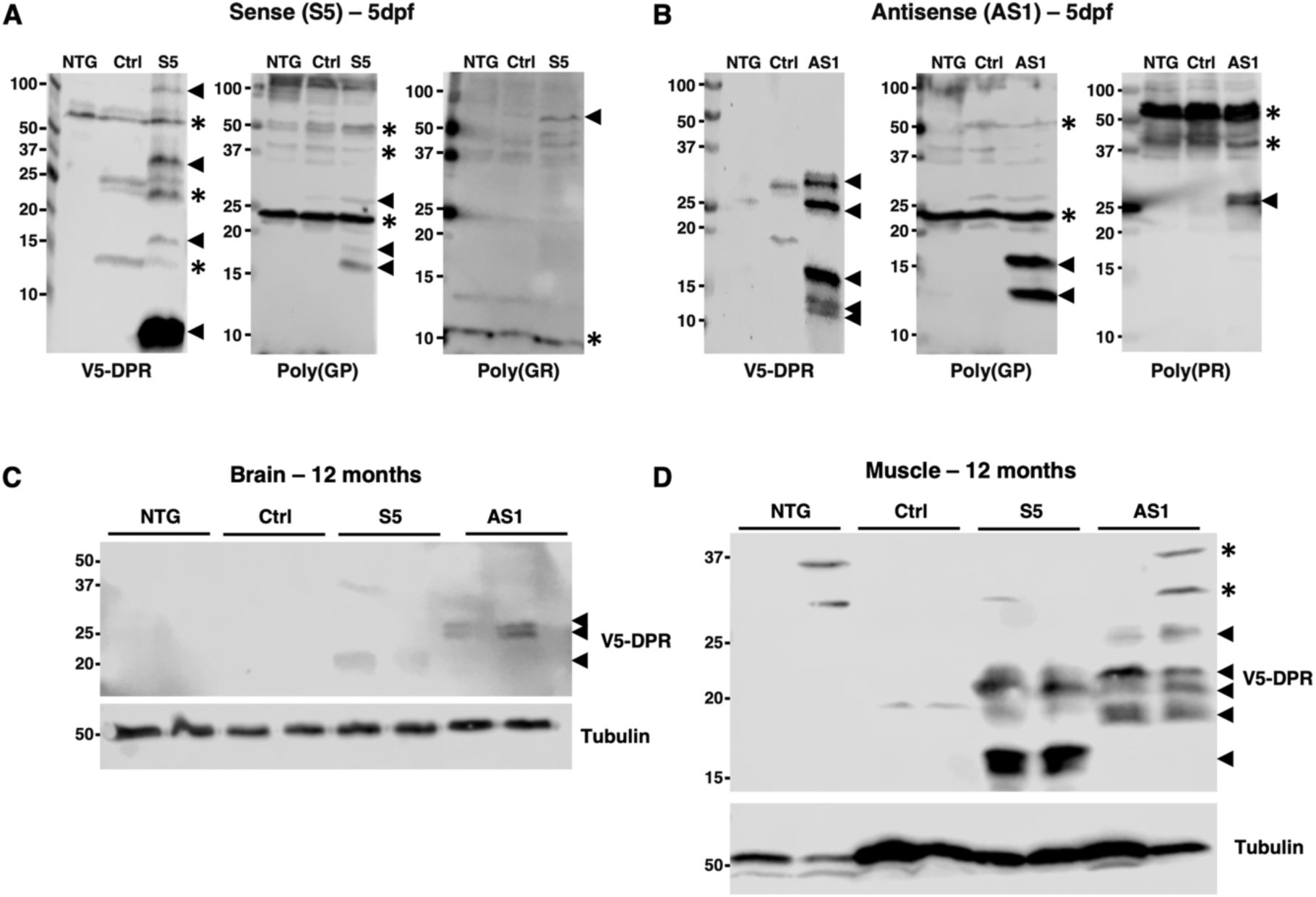
Embryonic and adult C9orf72-ALS/FTD zebrafish lines produce multiple RAN-translated DPR species. V5-DPRs and poly-repeats species GP, GR and PR in (a) sense (S5) and (b) anDsense (AS1) 5dpf embryonic lysates compared to non-transgenic (NTG) and 2mer (Ctrl). V5-DPR species detected in 12-month (c) brain and (d) muscle lysates of sense (S5) and anDsense (AS1) C9orf72-ALS/FTD zebrafish. Arrows indicate DPR bands which are proposed to be driven by RAN-translaDon. Asterisks indicate non-specific anDbody binding.

The presence of DPRs in embryonic 5 dpf embryos is a valuable tool for studying the mechanisms of DPR pathology and for screening tool compounds that may modulate this deleterious pathway. This said, C9orf72-mediated toxicity mainly occurs within the CNS and muscle tissue of ALS patients. Therefore, the next aim was to determine and characterize the presence of DPR species in adult zebrafish muscle and brain tissue lysates. V5-tagged DPR species were detected in both adult sense S5 and antisense AS1 zebrafish lines in both brain and muscle tissue lysates (Figures 3C-D). DPRs were observed at a higher molecular weight in the antisense AS1 line when compared to the sense S5 line in both brain and muscle tissue.

In summary, sense line S5 and antisense line AS1 produce multiple DPR species at 5 dpf and 12 months. Western blot analysis demonstrated that ribosomal slippage and frameshifting may be producing multiple DPR species above the predicted molecular weight. The locality of DPR expression replicates that seen in human C9orf72-ALS/FTD patients, further supporting the utility of this model as a screening tool for modulating DPR production in embryonic and adult zebrafish.

### Reduced swimming endurance in adult C9orf72-ALS/FTD zebrafish

Swimming endurance was tested to evaluate the neuromuscular integrity of adult transgenic zebrafish. Swim tunnel performance analysis showed that at both 6- and 12-month timepoints, swimming endurance was decreased in both sense S5 and antisense AS1 zebrafish, when compared to the transgenic 2mer (Ctrl) line. Quantification of endurance deterioration showed that swim tunnel performance significantly worsened between 6 and 12 months in both sense S5 and antisense AS1 transgenic zebrafish. This worsening was not observed in the transgenic 2mer (Ctrl) and therefore demonstrates the neurotoxic effects of RAN-mediated V5-DPR expression in the sense S5 and antisense AS1 transgenic lines (Figures 4A-C).

**Figure 4.**
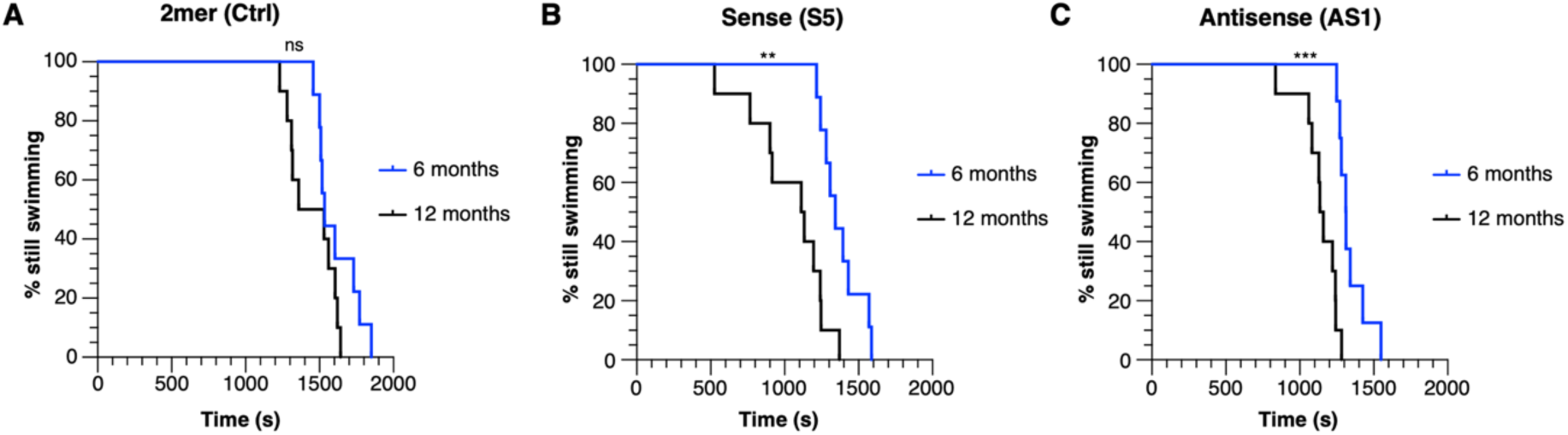
Adult C9orf72-ALS/FTD zebrafish show reduced swimming endurance. Kaplan-Meier analysis of swim tunnel performance in 6-month 12-month-old zebrafish. (a) 2mer (Ctrl) zebrafish show no significant decline in swimming endurance between 6- and 12-month timepoints. Both (b) sense (S5) and (c) antisense (AS1) zebrafish lines significantly decline in swimming endurance between 6- and 12-months old. N=9 2mer (Ctrl), N=10 sense (S5) and N=10 antisense (AS1). Log-rank (Mantel-Cox) test; *P < 0.05, **P < 0.01, ***P < 0.001.

In summary, deficits in swim tunnel performance are seen as early as 6-months-old and this loss of swimming endurance worsens up to the 12-month-old time point tested. The presence of adult-onset swimming endurance deficits provides an important phenotypic parameter to measure disease progression.

### Hypothermic cooling reduces DPR expression in C9orf72-ALS/FTD in vitro models

To build on this work, hypothermic conditioning and subsequent induction of the cold shock response was investigated as a mechanism to reduce RAN-translated DPR production. To test this, HEK293T, N2A cells and HeLa cells were transfected with the original RAN translation reporter constructs expressing 45 sense or 43 antisense repeats^17^. In all cell models, western blot analysis demonstrated that transfected cells show significant RAN-translated V5-DPR production in both sense and antisense cell lysates. No detectable V5-DPRs were observed in each of the control conditions. Hypothermic cooling to 32°C significantly reduced V5-DPR expression in all cell models in both sense and antisense C9orf72-ALS/FTD cell models when compared to normothermic (37°C) conditions (Figures 5A-D). Furthermore, immunostaining for cold shock proteins cold-inducible RNA-binding protein (CIRBP) and RNA-binding motif 3 (RBM3) were both significantly increased in hypothermia-treated cells, evidencing the activation of the cold-shock response (Figures 5E-G). Taken together, hypothermic cooling reduces RAN-translated V5-DPR expression in three separate mammalian cell lines which is linked to a significant increase in the cold shock response.

**Figure 5.**
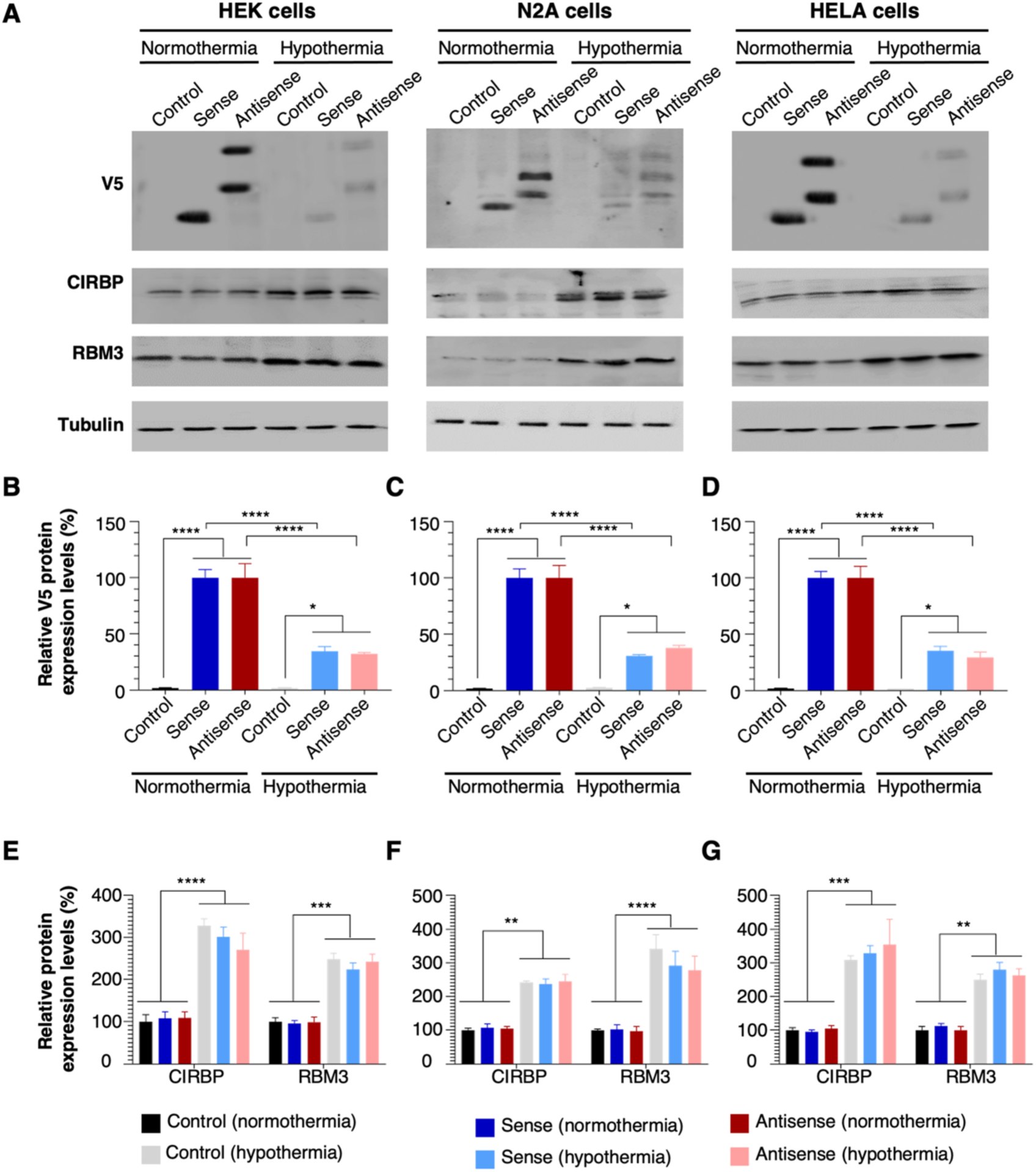
Hypothermic cooling reduces DPR expression in C9orf72-ALS/FTD cell models. (a) HEK293T, N2A and HeLa cells were transfected with control, sense and antisense constructs and protein lysates were collected following 48 h hypothermic cooling (32°C). Western blot analysis and quantification of (b-d) V5-DPRs and (e-g) cold shock proteins CIRBP and RBM3. Data presented as mean ± SD. One-way ANOVA; *P < 0.05, **P < 0.01, ***P < 0.001, ****P < 0.0001.

### Hypothermic-TTM reduces DPR expression in embryonic C9orf72-ALS/FTD zebrafish

To build on this, we further applied this hypothermic cooling protocol to embryonic transgenic C9orf72-ALS/FTD zebrafish to assess *in vivo* efficacy. Embryos were placed in normothermic (28°C) and hypothermic (22°C) conditions from 2 to 5 dpf. Western blot analysis of embryo lysates demonstrated a significant decrease in V5-DPR expression following hypothermic cooling in the transgenic sense S5 line (normothermia 100% ± 29.83 v hypothermia 18.6% ± 13.93, ***P = 0.0006, Figures 6A-B). Poly(GP) expression was also significantly decreased following hypothermic cooling of antisense AS1 transgenic embryos when compared to normothermic control embryos (normothermia 100% ± 33.54 v hypothermia 58.58% ± 23.16, *P = 0.0320; Figures 6E-F).

**Figure 6.**
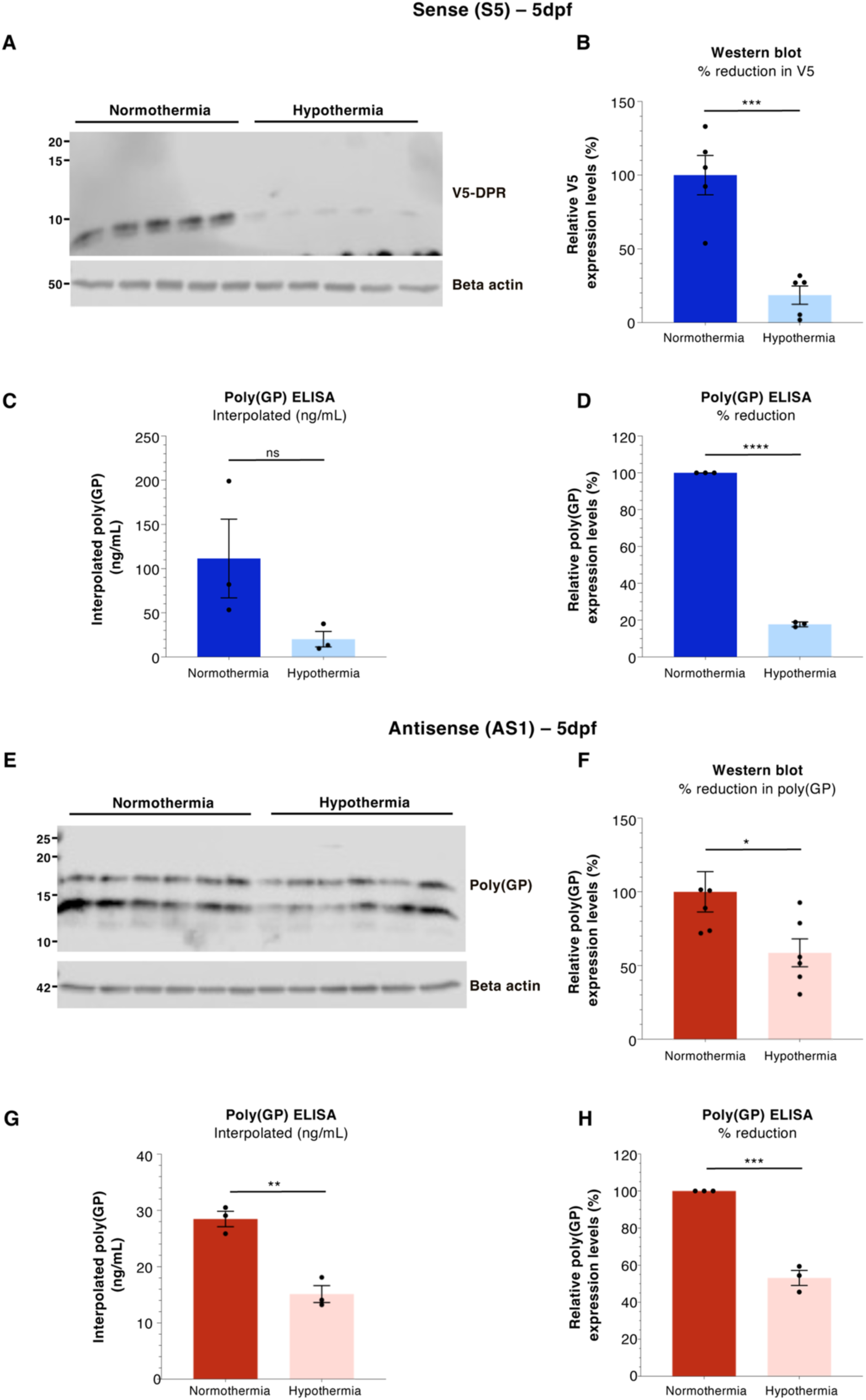
Hypothermic-TTM reduces DPR production in 5 dpf embryonic C9orf72-ALS/FTD zebrafish. Western blot and MSD poly(GP) ELISA analysis of 5 dpf embryonic lysates following 72 h hypothermic cooling. (a-b) anti-V5-DPR immunostaining and (c-d) MSD poly(GP) ELISA analysis of sense (S5) embryonic lysates. (e-f) anti-poly(GP) immunostaining and (g-h) MSD poly(GP) ELISA analysis of antisense (AS1) 5dpf embryonic lysates. MSD ELISA assay responses were interpolated using a GP(7) standard prior to normalisation. Western blot – N=5-6 clutches per treatment group; poly(GP) MSD ELISA – N=3 clutches per treatment group. Data presented as mean ± SD. Unpaired t-test, *P < 0.05, **P < 0.01, ***P < 0.001, ****P < 0.0001.

C9orf72-ALS/FTD sense and antisense embryonic lysates were further analyzed for poly(GP) expression using the Meso Scale Discovery (MSD) ELISA platform. Similar to the western blot analysis, MSD ELISA analysis showed a significant reduction in poly(GP) DPR species following hypothermic cooling when compared to their normothermic counterparts. This effect was seen in both sense S5 embryos (hypothermia 17.7% ± 1.32, ****P < 0.0001; Figures 6C-D) and antisense AS1 embryos (hypothermia 53.1% ± 7.00, ***P = 0.0003; Figures 6G-H). Taken together, this further shows the strong effect of hypothermic cooling on RAN-translation and subsequent DPR production.

## DISCUSSION

We describe a novel transgenic zebrafish RAN-translation model of C9orf72-ALS/FTD that expresses multiple RAN-translated DPR species in both embryonic and adult transgenic zebrafish. This recapitulates a major pathological cellular hallmark as well as the marked motor decline that is observed in C9orf72-ALS/FTD patients. Taken together, this model offers a novel pre-clinical research tool to further illuminate the pathological mechanisms of RAN-translation in the context of C9orf72-ALS/FTD, facilitating the screening and development of novel therapeutic approaches. We highlight this utility by demonstrating that reducing environmental temperature through hypothermic-TTM reduces RAN-translated DPR species in both C9orf72-ALS/FTD *in vitro* models as well as the C9orf72-ALS/FTD zebrafish models. This is the first study to demonstrate hypothermic-TTM as a viable therapeutic strategy to explore in C9orf72-ALS/FTD.

Multiple RAN-translated DPR species can be detected in sense S5 and antisense AS1 zebrafish embryos at 5 dpf. Incorporation of a V5 epitope tag in all three reading frames following the repeat expansion further facilitates higher throughput for screening changes in global RAN-translation *in vivo*. These pure repeat models recapitulates the HRE RAN-translation mechanisms that are observed in C9orf72-ALS/FTD patients^21^ and will facilitate further research to understand how RAN translation and the production of toxic DPR species may precipitate and/or exacerbate ALS progression, and therefore how these mechanisms may be targeted for therapeutic intervention.

### Zebrafish models of C9orf72-ALS/FTD

Zebrafish have emerged as a major pre-clinical research tool in recent years, particularly in neuroscience research studying complex brain disorders^22^. Zebrafish share 70% gene homology with humans and the structure of the zebrafish CNS has all the major mammalian brain domains and many of the same neurotransmitters^23^. Furthermore, the zebrafish motor system is largely similar to that of humans and many of the genes that are associated with familial ALS/FTD development risk are found in zebrafish, including C9orf72^24^.

A number of transient and stable ALS zebrafish models have previously been developed to recapitulate the cellular pathology and behavioural phenotypes that are observed in human C9orf72-ALS/FTD. A transient transgenic zebrafish model expressing 32 and 72 sense (GGGGCC) repeats exhibits increased apoptotic markers and DPR expression at 24 hpf when compared to transgenic embryos expressing only 8 sense repeats, where these markers were absent. This highlights the requirement of a minimum repeat number threshold to produce DPR species and induce cell death^25^. Other studies investigating motor performance and repeat expansion length found zebrafish with ≤ 10 repeat expansions exhibited no motor-axon defects or loss of motor performance, whereas transgenic lines containing 37, 70 or 90 repeats exhibited significant motor-axon abnormalities^26^. These studies are supported by our previous paper detailing motor decline and eventual early mortality in transgenic zebrafish stably expressing 89 hexanucleotide repeats^10^. In the present study, we observe no evident expression of RAN-translated DPR species or decline in motor performance in the transgenic control line expressing 2 sense repeats (2mer Ctrl). However, expression of either 45 sense (GGGGCC) or 39 antisense (CCCCGG) repeats results in the expression of multiple RAN translated DPR species as well as progressive motor performance decline, supporting previously published research.

Although this study focusses on the neurotoxic RAN-mediated DPR mechanisms of disease, characterisation of other C9orf72-ALS/FTD disease hallmarks has not yet been explored. Previously reported gain of function zebrafish and cell models of C9orf72-ALS/FTD have described motor neuron toxicity following expression of either sense^26, 25^ or antisense^10^ HREs, however TDP-43 mis-localization was not reported. This said, miRNA-based gene silencing of C9orf72 results in both TDP-43 mis-localization and motor neuron loss in a loss-of-function zebrafish model of C9orf72-ALS/FTD^27^. Further characterisation of motor neuron integrity and TDP-43 proteinopathy in adult CNS and peripheral tissues may illuminate further disease hallmarks in the RAN-translation C9orf72-ALS/FTD zebrafish models described.

### Hypothermic-TTM – cellular mechanisms

Targeted temperature management (TTM), previously termed therapeutic hypothermia, is the intervention to alter core body temperature to prevent fever, induce hypothermia or maintain normothermia. The neuroprotective effects of hypothermia have been explored in a number of pre-clinical and clinical studies targeting acute neurological disease. Through these studies, treatment protocols have been optimised and the mechanisms that underpin this neuroprotection have been interrogated. Although these mechanisms have not been fully elucidated, a number of neuroprotective pathways have been reported.

One of the first cold-shock proteins identified was RBM3^28^. Following hypothermic-TTM, RBM3 expression is increased through BDNF-TrkB signalling. This inhibits the action of phosphorylated ERK branch activation and is an important pathway in neuronal structural plasticity^29^. Furthermore, RBM3 expression localises to dendrites and increases local dendritic protein synthesis^30^ as well as global protein synthesis^31^ through binding of ribosomal subunits, specifically reticulon 3 (RTN3). RTN3 is involved in synaptic formation and neuronal plasticity^32^. *In vitro* studies have shown that RBM3 is substantially responsible for the neuroprotective effects of hypothermic-TTM. Using neuron-like PC12 cells, Chip *et al.* showed that blocking RBM3 through specific short interfering RNAs abolished the neuroprotective effects of cooling to 32°C. Furthermore, RBM3 overexpression reduced apoptotic markers in the absence of hypothermia^33^. The role of RBM3 has been further reported in animal models of neurodegeneration. Hypothermic-TTM significantly improves neurological outcomes in mouse models of Alzheimer’s disease and prion disease, however this neuroprotective capacity is reduced following lentiviral-mediated knockdown of RBM3. Furthermore, lentiviral-mediated overexpression promotes neuroprotection, with RBM3 over-expressing mice showing a marked increase in lifespan, later onset of neurodegeneration and improved behavioural outcomes when compared to knockdown mice^34^. Lastly, a recent paper identified that RBM3 pre-mRNA undergoes alternative splicing that is sensitive to changes in temperature. A previously uncharacterized exon (exon3a) is not included in the mRNA at 34°C, however exon3a-containing mRNA isoforms are present at 38°C. Furthermore, antisense oligonucleotide administration targeting exon3a results in a sustained increase in RBM3 expression as well as neuroprotection in a mouse model of prion disease^35^. These mechanisms could be further explored in the context of C9orf72-ALS/FTD.

CIRBP also plays a key role in the neuroprotective effects of hypothermic-TTM. CIRBP is an RNA-binding factor and expression is increased following hypothermia^36^. Mild hypothermia increases CIRBP expression in the cortex, hypothalamus and hippocampus^37^. *In vitro* studies using rat primary cortical neurons showed that CIRBP expression is protective against H_2_O_2_- induced apoptosis^38^. Experimental models of cerebral ischemia show evidence of a significant reduction in ROS and jugular NO release following mild hypothermia when compared to normothermic conditions^39^. Furthermore, using a rat model of TBI, the anti-apoptotic effects of mild hypothermia were abolished following CIRBP siRNA silencing^37^, indicating a significant neuroprotective role of CIRBP in hypothermic-TTM.

In this study, cooling C9orf72-ALS/FTD cell models to 32°C increased protein expression of both RBM3 and CIRBP and this is associated with a significant decrease in DPR expression. This potent effect of hypothermic-TTM on DPR expression was further observed in our C9orf72-ALS/FTD zebrafish models. The mechanisms that link hypothermic-TTM with a reduction in RAN-translated DPR species warrants further investigation. Previous studies have shown that cold temperatures enhance proteasome activity and reduce protein aggregation in *Caenorhabditis elegans* that express an ALS-related mutant variant of FUS protein^40^. Further experiments to discriminate between reduced DPR production and enhanced DPR clearance will elucidate the mechanistic pathways that influence RAN-translated DPR expression following hypothermic-TTM.

### Hypothermic TTM - clinical perspectives

Pre-clinical research efforts have demonstrated key pathways involved in efficacy outcomes following hypothermic-TTM. Progression of this research into randomized clinical trials has shed light on the utility of hypothermic-TTM in human disease. Hypothermic TTM has found most of its application in severe acute injury and is applied over a short period. The most widely cited and implemented use of hypothermic-TTM is in cardiac arrest (CA) patients. Following a large-scale multi-centre randomized control trial (RCT) the Hypothermia after Cardiac Arrest (HACA) Study Group found that mild hypothermia improves neurological outcome and reduces mortality in CA patients^41^. Hypothermic-TTM to a core body temperature of 32-34°C is now a recommended treatment protocol by the American Heart Association (AHA) for CA patients^42^.

Hypothermic-TTM has also been translated into the clinic for the treatment of hypoxic-ischaemic encephalopathy (HIE)^43, 44^. A recent meta-analysis of 28 RCTs found that hypothermic-TTM reduced the risk of mortality in neonates with moderate to severe HIE, concluding that healthcare professionals should consider this as a treatment for neonates with HIE^44^.

Hypothermic-TTM has also been explored in the context of ischaemic stroke. A meta-analysis conducted by van der Worp *et al.* concluded that therapeutic hypothermia improves outcomes by approximately one third in animal models of cerebral ischaemia, in conditions that are achievable in a clinical setting^45^. This clear evidence warranted further investigation in human patients, however to date, this pre-clinical research has not been successfully translated into an effective therapy for ischaemic stroke patients. A meta-analysis comprising twelve studies found no additional benefit in hypothermic-TTM ischaemic stroke patients. Furthermore, treated patients experienced higher adverse events, which may be due to the presence of patient co-morbidities increasing the risk of adverse event development. The influence of comorbidities on treatment outcome needs to be addressed in future studies^46^. Research efforts to standardise treatment protocols and outcome measurements as well as to further understand the influence of age, gender and comorbidities on these outcomes are needed to translate pre-clinical research efforts into a safe and effective treatment option^47, 48^.

Efforts have been made to improve therapeutic hypothermic-TTM protocols, including the use of hydrogel-coated water-circulating transfer pads^49^ or cold fluid intravenous infusions^50^ to improve upon more conventional cooling systems. Supplementary treatment during periods of hypothermic cooling may also reduce the risk of adverse event development. These alternative and refined cooling approaches, alongside supplementary treatment approaches, allow more controlled hypothermia induction and maintenance, which may provide more safe and practical options for prolonged use in future clinical studies^51^. To what extent these different treatment modalities influence experimental and clinical outcomes needs to be defined in disease-specific contexts, thus careful study-design is paramount to ensure patient safety and determine clinically efficacy.

## Conclusions

The transgenic model described in this paper provides a medium throughput *in vivo* research tool to further investigate the toxic role of RAN-translation in C9orf72-ALS/FTD, and further understand the mechanisms that underpin any neuroprotective strategies. In comparison to current murine models of C9orf72-ALS/FTD, the zebrafish model described offers a higher throughput pre-clinical model that could be utilized to assess both target efficacy and motor improvement in a non-mammalian system.

Future work aims to further explore the mechanisms of hypothermic-TTM by utilising the C9orf72-ALS/FTD cell models and the RAN-translation *zebrafish* models described. This will allow us to further understand how hypothermia influences C9orf72-ALS/FTD tissue pathology and motor performance. Additionally, combining hypothermic-TTM protocols with other neuroprotective strategies may provide synergistic effects, as highlighted by previous studies of neurodegenerative disease^52^. The ultimate aim of this research is to eventually translate these pre-clinical data into an effective therapy for C9orf72-ALS/FTD. The substantial array of safety and efficacy data that has been published to date clearly demonstrates the potential benefit of hypothermic-TTM in neurological disease.

## ACKNOWLEDGEMENTS

We acknowledge Dr Andrew Grierson for useful discussions and comments on the manuscript. T.R., P.J.S. and G.M.H. acknowledge MND Association research grant Apr17/854-791. G.M.H. further acknowledges support from the Medical Research Council (MRC) New Investigator research grant MR/R024162/1 and the Biotechnology and Biological Sciences Research Council (BBSRC) grant BB/S005277/1. The authors declare having no conflict of interest.

## AUTHOR CONTRIBUTIONS

T.R., G.M.H. and P.J.S funded, conceived, designed and supervised the study, D.J.B, A.M., O.A. and L.M.C. acquired and analysed the data, D.J.B drafted the manuscript and figures. All authors edited and approved the final manuscript.

